# Regulation of the innate immune response in human neurons by ICP34.5 maintains herpes simplex virus 1 latency

**DOI:** 10.1101/2025.04.04.647253

**Authors:** Paige N. Canova, Sarah Katzenell, Stacey Cerón, Audra J. Charron, Jean M. Pesola, Hyung Suk Oh, Donald M. Coen, David M. Knipe, David A. Leib

## Abstract

Herpes simplex virus 1 (HSV-1) establishes latent infections in sensory neurons, from which HSV sporadically reactivates due to external stress and other stimuli. Latency and reactivation are studied through *in vivo* models in a variety of hosts, as well as *in vitro* models using primary neurons, and neurons derived from pluripotent stem cells (iPSCs). These systems behave disparately, but the reasons remain unknown. The interferon (IFN)-based neuronal innate immune response is critical in controlling HSV-1 replication and HSV-1 counters these responses in part through infected-cell protein 34.5 (ICP34.5). ICP34.5 also promotes neurovirulence by preventing host translational shutoff and interfering with host cell autophagy. Here we demonstrate in a human iPSC neuronal model that sustaining host translation is the key activity of ICP34.5 for enhancement of reactivation. Specifically, our data shows that ICP34.5 was key for maintenance of HSV-1 latency. While interaction of ICP34.5 with the autophagy regulator Beclin 1 was important for maintaining latency, this was not due to modulation of bulk autophagy. Our work from primary mouse neurons suggested that the major effect of ICP34.5 on latency maintenance occurs in an IRF3/7-dependent manner. Notably, the role of ICP34.5 in regulating latency and reactivation differs between neurons derived from human iPSCs (iNeurons) and primary mouse trigeminal (TG) neurons. This highlights the importance of selecting an appropriate neuronal model and validating experimental outcomes in multiple models.

## Introduction

Herpes simplex virus 1 (HSV-1) is a double-stranded DNA virus of humans that infects ∼70% of the global population[1]. Initial infection of HSV-1 occurs at mucosal membranes followed by spread to innervating sensory neurons. Once internalized capsids reach the neuronal cell body through retrograde transport, the virus establishes a lifelong latent infection that can reactivate to produce infectious virions in response to stimuli such as UV irradiation, stresses, or fever. Following reactivation, new virions travel within axons in an anterograde direction to the initial site of infection where they are released and replicate in epithelial cells, causing mucosal lesions. During viral infections, the innate immune system is activated through pattern recognition receptors (PRRs), sentinel proteins that detect pathogen proteins and nucleic acids. PRR-triggered immune induction can activate TANK-binding kinase 1 (TBK1) that in turn phosphorylates interferon regulatory factor 3 (IRF3). Phosphorylated IRF3 then translocates to the nucleus and promotes expression of type I interferons (IFN). Type I IFNs (IFN-α and IFN-β) function in an autocrine and paracrine fashion to initiate and amplify an antiviral state in infected and naïve cells[2]. A central mechanism in the IFN response is the shutdown of protein translation, limiting viral protein expression and thus precluding virion assembly. This is mediated largely by the double-stranded RNA-dependent protein kinase R (PKR), which phosphorylates and thereby inactivates the key protein translation factor eIF2α, resulting in translational shutoff and the antiviral state of the cell[3].

While IFN is essential in protecting neurons during an HSV infection, prolonged exposure to high levels can be detrimental to cell survival[4,5]. It is therefore critical that neurons moderate the immune response, and they do so in part through autophagy, which maintains cellular homeostasis through the degradation and recycling of cytoplasmic components or intracellular pathogens[6]. Autophagy is initiated in response to a cellular stress such as starvation or infection, which causes inhibition of the negative regulator mammalian target of rapamycin complex 1 (mTORC1)[7]. This in turn allows activation of the initiation UNC-51-like kinases (ULK) complex[8]. This complex then activates the class III PI3K complex comprising of Vps34, Beclin 1, p150, and Atg14L[9,10]. During the elongation and maturation stages of autophagy, microtubule-associated protein 1 light chain 3 (LC3) is conjugated with phosphatidylethanolamine (PE). PE-conjugated LC3 (LC3-II) is incorporated into the forming phagophore membrane[11]. Cellular or exogenous components are marked for degradation by autophagy through ubiquitination by autophagic adaptor proteins such as p62/SQSTM1. Once cargo is destined for degradation, p62 transports and anchors the cargo to the autophagosome through interaction with LC3-II[12]. Following autophagosome formation, fusion with lysosomes results in the degradation of contents and inner membrane by lysosomal hydrolase[13].

Autophagy can activate the cell-intrinsic immune system through trafficking events[14], creating a feedback loop in which immune effectors, notably PKR and phosphorylated eIF2α, in turn promote autophagy[15,16]. Autophagy can also negatively regulate the immune response through p62-mediated degradation of stimulator of IFN genes (STING), an adaptor protein upstream of TBK1[17]. HSV-1 encodes numerous genes to evade or manipulate the immune system and the autophagy pathway. The multifunctional infected-cell protein 34.5 (ICP34.5), in particular, is a major antagonist of the antiviral and autophagy response[18]. ICP34.5 acts throughout the lytic viral life cycle, and is critical for efficient replication and neurovirulence of HSV-1[19]. ICP34.5 contains a domain that binds to TBK1, causing TBK1 degradation and decreased IRF3 and IFN expression[20]. A central function of ICP34.5 counters the action of PKR on protein translation, through its recruitment of protein phosphatase 1-alpha (PP1α), ICP34.5 triggers the dephosphorylation of eIF2α allowing viral protein synthesis to proceed and limiting autophagy[21-23]. ICP34.5 also inhibits autophagy through its binding to Beclin 1, a protein required for the class III PI3K complex during initiation[19]. The Beclin 1 binding domain (BBD) of ICP34.5 is critical for HSV-1 pathogenesis and regulation of autophagosome formation in primary sympathetic neurons[19].

Our current understanding of the role of ICP34.5 in regulating autophagy and IFN, and how that regulation subsequently affects the viral lifecycle, has primarily been acquired with mice *in* vivo, and primary mouse neurons and fibroblasts *in vitro*. However, given that HSV has evolved to evade human immune systems, there are some key differences between mice and human immunity such that HSV infections are more effectively controlled in mice[24]. We therefore set out to examine the role of ICP34.5 in regulating the innate, cell-intrinsic response in a human neuron model. We were especially interested in determining how ICP34.5 impacts latency establishment and maintenance in human neurons, and uncovering any mechanistic differences in the IFN/ICP34.5 axis between human and mouse neurons. Using human sensory neurons (iNeurons) derived from induced pluripotent stem cells (iPSCs), we showed that the most impactful role of ICP34.5 in latency and reactivation is regulation of the IFN response, rather than modulation of autophagy. We also show that while ICP34.5 is not required to establish latency, it is important for maintaining latent infection in iNeurons. In contrast, in mouse neurons, ICP34.5 is critical for induced reactivation and the host IFN response is pivotal for the maintenance of latency in the presence or absence of ICP34.5. The data support the idea that the choice of neuronal model is a key driver of experimental outcomes when determining roles for immunomodulatory functions in a system with the complexity of HSV latency and reactivation.

## Results

### A NEUROG3-based differentiation pipeline generates mature neurons that support latency and reactivation

In order to study HSV in a human neuron model, we generated neurons from the human iPSC line iNGN3, which overexpresses neurogenin-3 (NGN3) in the presence of doxycycline[25]. NGN3 is a member of a family of transcription factors that affect the commitment of progenitors to neurons by promoting neuronal subtype-specific gene expression[26]. NGN3 overexpression in the iPSCs induces differentiation into a neuron-like morphology within 4 days[27], significantly shorter than typical iPSC differentiation protocols that can take several weeks. A combination of inhibitors and growth factors were used to promote differentiation towards a human sensory neuron phenotype as described previously[28-31, Oh *et al*. Manuscript under review] (Figure 1A). The differentiating cells gradually transition in culture morphology and cell shape (Figure 1B; Supplemental 1) and express canonical markers of mature neurons including Tuj-1 and NeuN, while lacking expression of the proliferation marker Ki-67 (Figure 1C). iNGN3 cells, on the other hand, did not express NeuN, but had Ki-67 proliferation expression as expected (Figure 1C). Therefore, this rapid differentiation pipeline provides a renewable source of iNeurons, mature human sensory neuron-like monocultures, to model HSV-1 infections free from the complexities inherent in mixed or poorly defined cultures.

**Figure 1:**
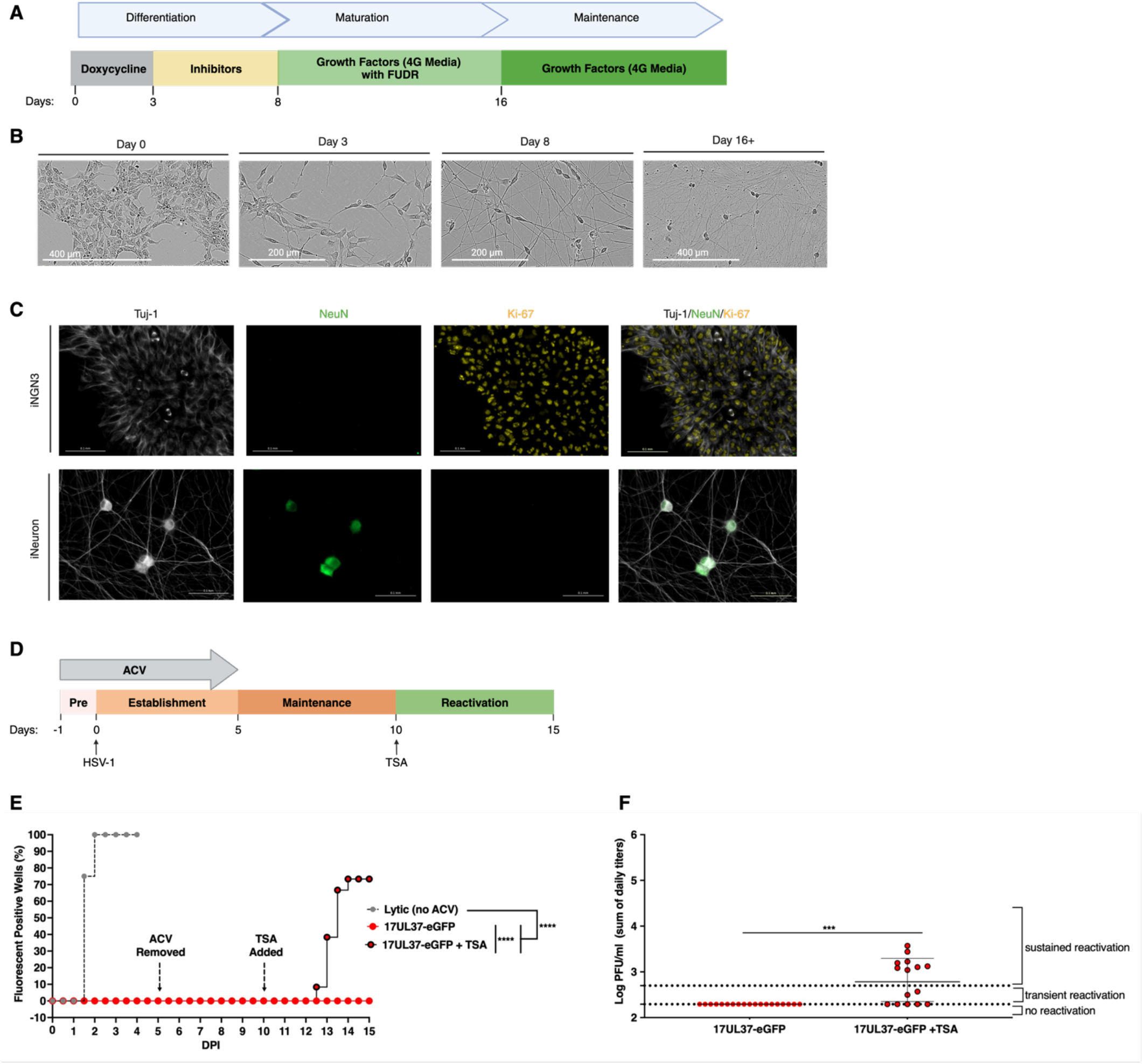
Differentiation and characterization of iNGN3s into mature sensory neurons as model cells for establishment of HSV-1 quiescence. (**A**) Timeline of the iNGN3 differentiation into mature neurons (iNeurons). (**B**) Representative phase-contrast images of 0-16 days post doxycycline-induced differentiation of iNGN3 cells. (**C**) Representative immunofluorescence images of undifferentiated iNGN3 cells and 16 days post differentiated iNeurons stained with antibodies for neuronal markers Tuj-1 (white) and NeuN (green), and proliferation marker Ki-67 (yellow). Scale bar = 0.1 mm (**D**) Timeline of establishment, maintenance, and reactivation of latency. Acyclovir (ACV) was added to the culture one day prior to infection and maintained until 5 days post infection (DPI). iNeurons were infected on day 0 with HSV-1 strain 17 UL37-eGFP (17UL37-eGFP) at an MOI of 1. Reactivation was induced 10DPI with trichostatin A (TSA) which was removed 11DPI. (**E**) HSV-1 latency and reactivation monitored through fluorescence (UL37-eGFP expression) from 0-15DPI. Graph depicts the average percentage of wells expressing eGFP from at least 4 wells per condition in at least 2 separate experiments. (**F**) The sum of the viral titers from 10-15DPI plotted as the mean of at least 4 wells from >2 experiments. The dotted lines represent the limit of detection and the threshold for sustained reactivation. Data analyzed by Log-rank (Mantel-Cox) tests (**E**) or Welch’s t-test (**F**). ***, p ≤ 0.001; ****, p ≤ 0.0001.

Almost all cultured cells exhibit susceptibility to HSV lytic infection, but a more unusual and desirable feature is the support of viral latency and subsequent reactivation. iNeurons support lytic HSV-1 infection similar to other *in vitro* cell models[32], yet are untested as a model of HSV-1 latency and reactivation. Differentiated iNeurons were infected with HSV-1 under conditions that have been shown to promote viral latency in other systems[33]. To prevent lytic infection, iNeurons were treated with acyclovir (ACV), causing termination of viral DNA replication and thus precluding the expression of late lytic genes. ACV was added to iNeurons one day prior to infection, and remained in the media for 5 days post infection (DPI) (Figure 1D). We utilized 17UL37-eGFP[34], an HSV-1 strain 17 recombinant virus containing eGFP fused to the late lytic gene UL37, in conjunction with live cell imaging to monitor viral protein expression over two weeks of HSV-1 quiscence/latency. During acute/lytic infection in the absence of ACV, fluorescent protein expression was detected with Incucyte live-cell imaging in every culture well within 48 hours post infection (HPI) (Figure 1E, grey symbols). In contrast, during quiescent/latent infection established in the presence of ACV at a viral multiplicity of infection (MOI) of 1, UL37-eGFP was undetectable by 48HPI and beyond, even following the removal of ACV from culture media (Figure 1E, red symbols). To test whether latent/quiescent virus could reactivate, the histone deacetylase inhibitor trichostatin A (TSA) was added 10DPI. TSA promotes chromatin opening and results in access of transcription factors to lytic gene promoters. Starting 2 days after TSA treatment, many wells in the infected cohort became fluorescent, indicating onset of lytic UL37-eGFP expression and reactivation (Figure 1E, red and black symbols, and Supplemental 2), which correlated with an increase in viral particles (Figure 1F). In contrast, wells containing the infected iNeurons untreated by TSA were uniformly non-fluorescent, indicating quiescence out to 15DPI (Figure 1E and 1F, red symbols). These results validate iNeurons as a suitable *in vitro* model with which to study HSV-1 quiescence/latency and reactivation.

### The interaction of HSV-1 with Beclin 1 is required for maintenance of latency in iNeurons

Beta- and gammaherpeviruses utilize autophagy to modulate latency[35]. During lytic infection, HSV-1 ICP34.5 inhibits autophagy through interaction with Beclin 1; likely preventing phagophore formation through limiting Beclin 1 availability to be incorporated in the PI3K complex (Figure 2A)[19]. To determine whether this interaction is important for lytic HSV-1 replication in iNeurons, we used a recombinant virus (ΔBBD) in which the Beclin 1 binding domain was deleted from ICP34.5. As ICP34.5 is a multi-domain protein, we also tested a null virus lacking ICP34.5 (Δ34.5) to measure the impact of eliminating all functional domains critical for viral replication and host defense (Figure 2B). During acute infection of iNeurons, Δ34.5 produced significantly lower titers than wildtype (WT) HSV-1, while ΔBBD produced titers similar to that of WT (Figure 2C). Therefore, in iNeurons ICP34.5 is required for maximal viral replication, though the discrete interaction between ICP34.5 and Beclin 1 does not promote lytic viral production, consistent with previous results[36]. As ICP34.5 has domains that support viral replication, decreased viral production by Δ34.5 may result from dampened viral gene expression in iNeurons. Thus, using qRT-PCR we measured expression of the immediate early gene ICP27, the early genes ICP6 and ICP8, and the late gene gC. Infection with Δ34.5 resulted in comparable viral gene expression of all three temporal classes compared to that of WT or ΔBBD infection, with any observed differences not being statistically significant (Figure 2D). Additionally, protein expression detected by fluorescent reporter-tagged viruses was comparable between WT, ΔBBD, and Δ34.5 infected iNeurons during an acute infection (Figure 2E). Therefore, decreased viral production by iNeurons infected with Δ34.5 was not due to a global transcriptional down-regulation of HSV gene expression.

**Figure 2:**
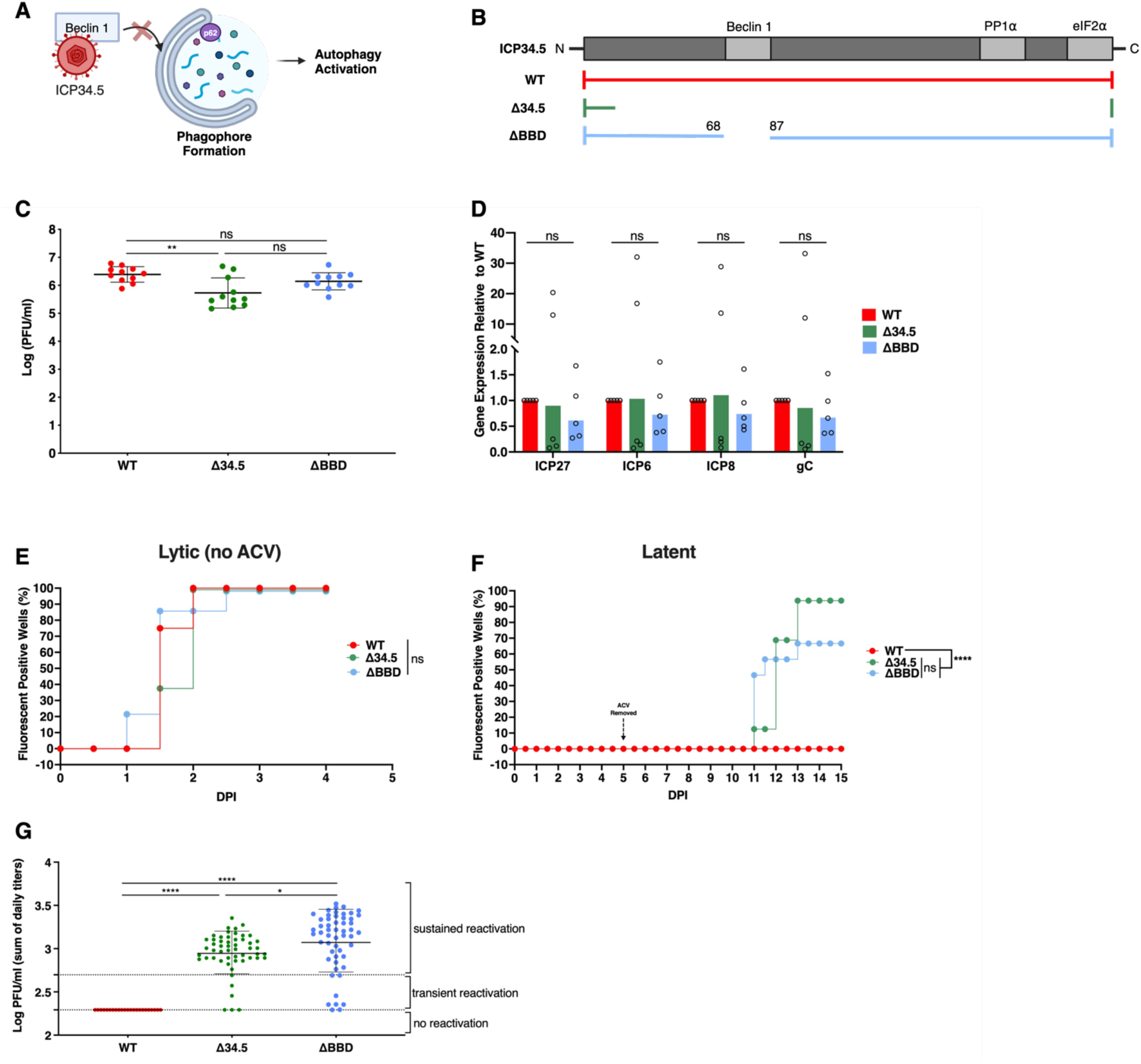
ICP34.5 interaction with Beclin1 is critical for latency maintenance in iNeurons. (**A**) Graphic depicting the likely interaction of HSV-1 ICP34.5 inhibiting autophagy phagophore initiation through its interaction with Beclin 1. (**B**) Map of the functional domains and recombinant viruses engineered into the ICP34.5 open reading frame of HSV-1 strain 17. (**C**) Viral titers at 24 hours post infection (HPI) of iNeurons infected with HSV-1 recombinant viruses at an MOI of 1. Each data point represents an individual well from three independent experiments. (**D**) RNA expression measured by qRT-PCR of ICP27, ICP6, ICP8, and gC viral gene expression by **Δ**34.5 and ΔBBD normalized to WT. iNeurons were infected at an MOI of 1, and collected 24 HPI. Samples were pooled from 3 wells from five independent experiments. Data from each independent experiment is depicted by open circles, and the bar depicts the geometric mean. (**E**) Graph depicting percentage of wells becoming fluorescent during lytic infection from 4-5 wells per condition in 2-3 separate experiments. (**F**) Percentage of wells becoming fluorescent during latent infection leading to spontaneous reactivation (no TSA treatment) of HSV-1 strain 17 UL37-eGFP (WT), ΔBBD mCherry, or Δ34.5 UL37-eGFP from 4-5 wells per condition in 2-3 separate experiments. (**G**) The sum of the viral titers from cultures shown in panel F (10-15 DPI) plotted as the mean of at least 4 wells from >2 experiments. The dotted lines represent the limit of detection and the threshold for sustained reactivation. Data analyzed by one-way ANOVA (**C** and **D**) with Šídák posttest, two-way ANOVA (**G**) with Šídák posttest, or Log-rank Mantel-Cox tests (**E** and **F**). ns = not significant; *, p ≤ 0.05; **, p ≤ 0.01; ****, p ≤ 0.0001. Error bars represent standard deviation from the geometric mean.

Since a defining characteristic of latency is suppression or absence of viral gene expression, the trend of increased lytic gene expression in Δ34.5 infection could indicate that ICP34.5 has a role in regulating latency. To test this iNeurons were latently infected with fluorescent reporter-tagged WT, ΔBBD, or Δ34.5 viruses and spontaneous reactivation from latency was monitored through fluorescent reporter gene detection (Figure 2F). All three viruses are competent to establish latency/quiescence in the presence of ACV as indicated by non-fluorescence in all wells from 0-10DPI (Figure 2F). This signifies that ICP34.5 is not essential for latency/quiescence establishment in iNeurons. However, 6 days after the removal of ACV from culture media, an increasing number of wells infected with ΔBBD and Δ34.5 became spontaneously fluorescent (Figure 2F, blue and green symbols), while wells infected with WT virus remained non-fluorescent (Figure 2F, red symbols). Fluorescent reporter expression correlated directly with virus production 10-15DPI as determined by plaque assay, ΔBBD- and Δ34.5-infected iNeurons had viral titers characteristic of sustained reactivation, while WT-infected iNeurons had no detectable viral production (Figure 2G). This finding suggests that ICP34.5, specifically the Beclin 1-binding domain of ICP34.5, plays a role in the maintenance of latency in iNeurons.

### Selective autophagy in iNeurons is unimpacted by HSV-1 infection and autophagy-modulating compounds

The interaction of ICP34.5 with Beclin 1, an important protein in autophagosome initiation, led us to hypothesize that autophagy regulation by the BBD promotes latency. However, it was important to first determine whether autophagy is impacted during infection by WT, ΔBBD, or Δ34.5 viruses. The selective autophagy adaptor protein p62 is consumed during autophagy, therefore the concentration of p62 is a convenient metric for autophagic flux. We expected WT virus infection to cause reduced levels of autophagy by virtue of interaction between Beclin 1 ICP34.5 (Figure 3A), as first demonstrated in primary mouse sympathetic neurons[19]. Conversely, we expected increased or unchanged levels of autophagy following infection with ΔBBD and Δ34.5. However, lytic infection with WT, ΔBBD, or Δ34.5 viruses had no impact on p62 levels in either undifferentiated iNGN3 cells or iNeurons compared to mock-infected control cells (Figure 3B). The unexpected resistance of selective autophagy to manipulation by ICP34.5 BBD-containing WT virus raised the possibility that iNGN3 cells are resistant to modulation of autophagy. To test this, we treated uninfected cells with the autophagy modulators LY294002, PIK-III, or rapamycin (Figure 3C). All three compounds intersect the autophagy pathway at the initiation stage, LY294002 and rapamycin activating autophagy and PIK-III inhibiting autophagy. Although rapamycin caused a slight increase in autophagy in iNGN3 cells (as measured by p62), only LY294002 treatment caused a significant increase in autophagy (Figure 3D). In iNeurons, unexpectedly, neither rapamycin nor LY294002 impacted p62 levels (Figure 3D). The VPS34 inhibitor PIK-III likely functions similarly to the BBD of ICP34.5 by inhibiting formation of the protein complex necessary to form a phagophore (Figure 3C). PIK-III treatment similarly affected iNGN3 cells and iNeurons, causing only a modest decrease in autophagy (Figure 3D). The resistance of iNeurons to potent autophagy modulators has precedent in primary neurons[37], where it is thought that autophagy is tightly regulated as a homeostatic mechanism[38]. In summary, HSV-1 infection, regardless of ICP34.5 status, failed to significantly alter autophagy in iNGN3 cells or iNeurons.

**Figure 3:**
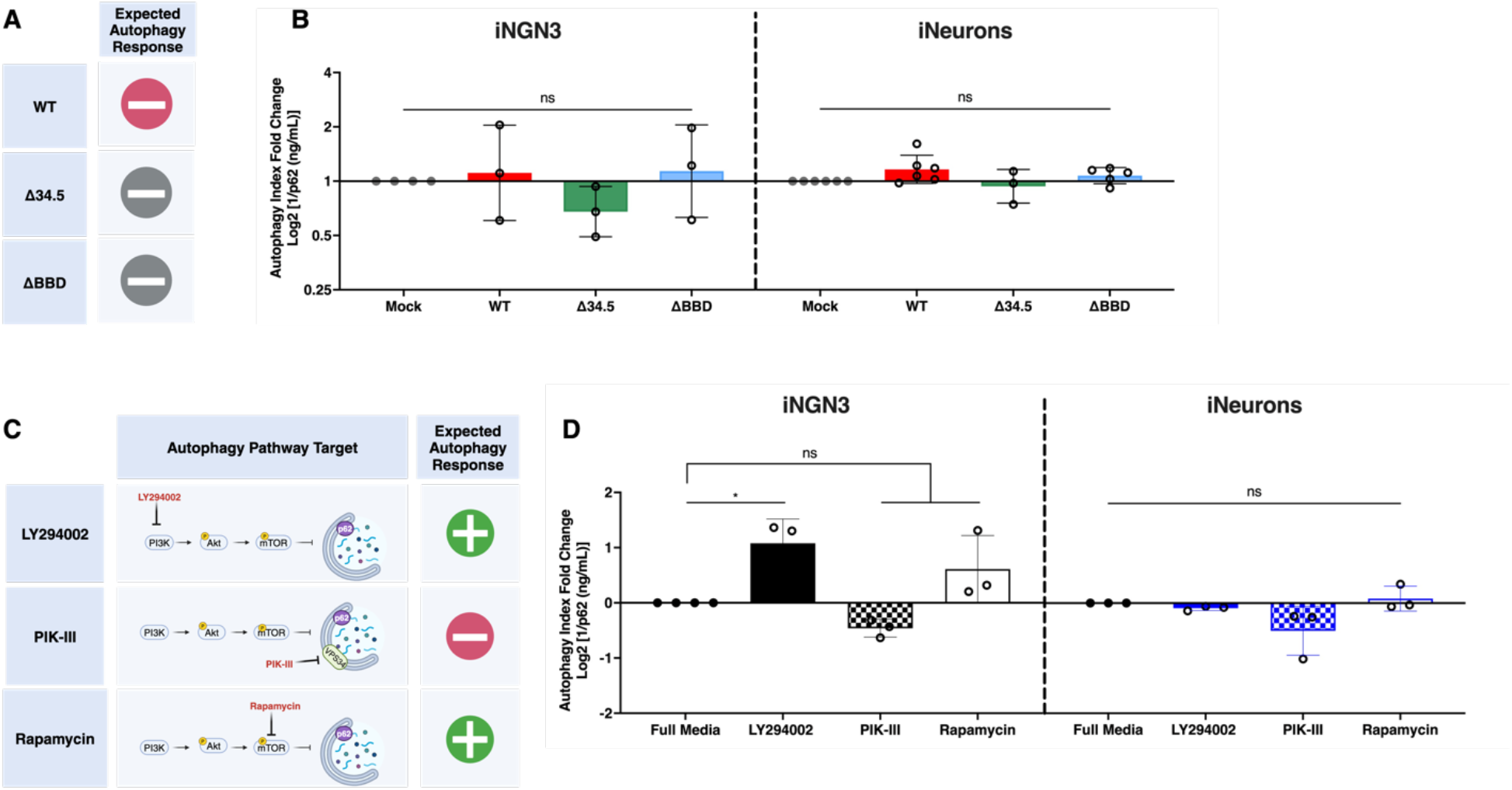
HSV-1 infection and autophagy-modulating drugs do not cause significant changes in p62 protein levels of iNGN3 cells or iNeurons. (**A**) Table depicting expected effect on autophagy during infection with HSV-1. Red circle with a minus sign depicts autophagy inhibition, and gray circle with a minus sign depicts minor to undetectable changes. (**B**) Graphs of p62 ELISAs of extracts from infected iNGN3 and iNeurons. Autophagy index represents the inverse of p62 protein concentration of iNGN3 cells and iNeurons relative to mock-infected cells. (**C**) Table of autophagy modulating treatments, their interactions in the pathways (red font) and the expected effects on the autophagy pathway. Green circle with a plus sign depicts autophagy induction, and red circle with a minus sign depicts autophagy inhibition. (**D**) Graphs of p62 ELISAs of extracts from treated iNGN3s and iNeurons. The autophagy index of iNGN3 cells and iNeurons are shown following treatment with LY294002, PIK-III, or Rapamycin for 24 hours. Data analyzed by one-way ANOVA (**B** and **D**) with Šídák posttest. ns = not significant; *, p ≤ 0.05. Bars represent the mean of three wells from three independent experiments (shown as a circle) and displayed as the geometric mean with error bars representing the SD.

### Autophagy modulators do not influence viral production

Autophagy, as judged by p62 levels, in iNGN3 cells and iNeurons was largely unchanged in response to HSV-1 infection or chemical modulators. We wished, however, to elucidate whether autophagy regulation impacted the lifecycle of HSV-1 independently of p62. iNGN3 cells and iNeurons were infected with a low MOI of HSV-1 in the presence of LY294002 or PIK-III for 24 hours and viral replication quantified (Figure 4A). LY294002 treatment, which promoted autophagy in uninfected iNGN3 cells decreased viral titers in WT- and Δ34.5-infected iNGN3 cells, and slightly decreased replication in ΔBBD-infected iNGN3 cells but to a non-significant extent (Figure 4B). PIK-III, which inhibited autophagy in uninfected iNGN3 cells, diminished viral replication in WT-, Δ34.5-, and ΔBBD-infected iNGN3 cells (Figure 4B). Remarkably, viral replication in WT-, Δ34.5-, and ΔBBD-infected iNeurons was unimpacted by addition of any autophagy modulator tested (Figure 4C). These results signify that the strict regulation of autophagy in iNeurons buffers pharmacological or viral modulation of this pathway.

**Figure 4:**
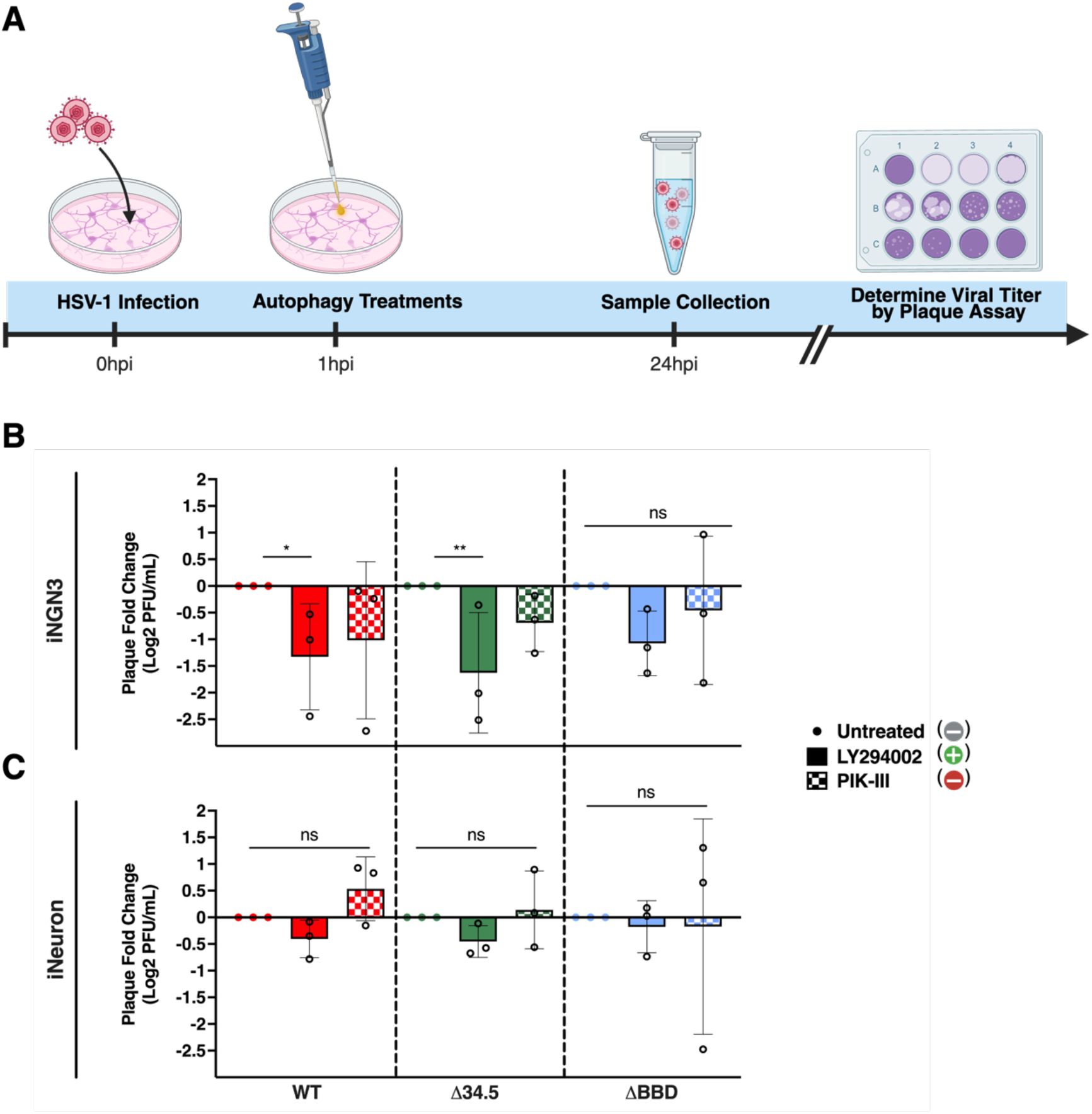
Viral production in iNGN3s and iNeurons are not impacted by autophagy modulating drug treatments. (**A**) Timeline of infection with autophagy treatments. Cells were infected with HSV-1 recombinant viruses at an MOI of 1, treated with either LY294002 or PIK-III, and collected 24 hours post infection (HPI). Viral titers from infected iNGN3s (**B**) and iNeurons (**C**) shown as a fold change relative to their untreated controls. Expected autophagy responses are indicated by colored circles in the legend. Each bar represents the mean of three wells repeated in three independent experiments and displayed as the geometric mean with the error bars representing the SD. Data analyzed by one-way ANOVA (**B** and **C**) with Šídák posttest. ns = not significant; *, p ≤ 0.05; **, p ≤ 0.01.

### The PP1α domain of ICP34.5 is essential for countering the IFN response in iNeurons

We next wished to address the hypothesis that, as in other cells and systems[4,39-41], ICP34.5 modulates IFN-driven immunity in iNeurons. In response to neuronal infection by HSV-1 *in vivo*, IFN is released by surrounding microglia and astrocytes, and curbs further neuronal infection[39,42,43]. We questioned whether similarly, IFN treatment could confer resistance of iNeurons to HSV-1 infection. To establish experimental parameters, we used primary human foreskin fibroblasts (HFFs) in which ICP34.5-deficient HSV-1 strains are sensitive to IFN-β[2]. HFFs were pre-treated with IFN-β for 18 hours, then infected with HSV-1, or with prototypical IFN-sensitive RNA viruses [44] namely vesicular stomatitis (VSV) and yellow fever (YFV) (Figure 5A). In IFN-β-treated HFFs, VSV replication was decreased by >40,000-fold, and YFV replication was reduced ∼700-fold (Figure 5B). As expected, replication of the HSV-1 variants in HFFs were also impacted by IFN pretreatment with WT, ΔPP1α, and ΔBBD having a ∼70-fold reduction. Notably, the null Δ34.5 virus was more drastically reduced with a >20,000-fold reduction, comparable to the sensitivity observed for VSV, the prototypical IFN-sensitive virus. IFN-β pre-treatment of iNGN3 cells, conversely, had no significant effect on the titers of any virus tested (Figure 5C). This unresponsiveness to IFN, and notably following infections with the highly IFN-sensitive VSV, was consistent with the observation that iPSCs lack robust responses to type I IFNs[45].

**Figure 5:**
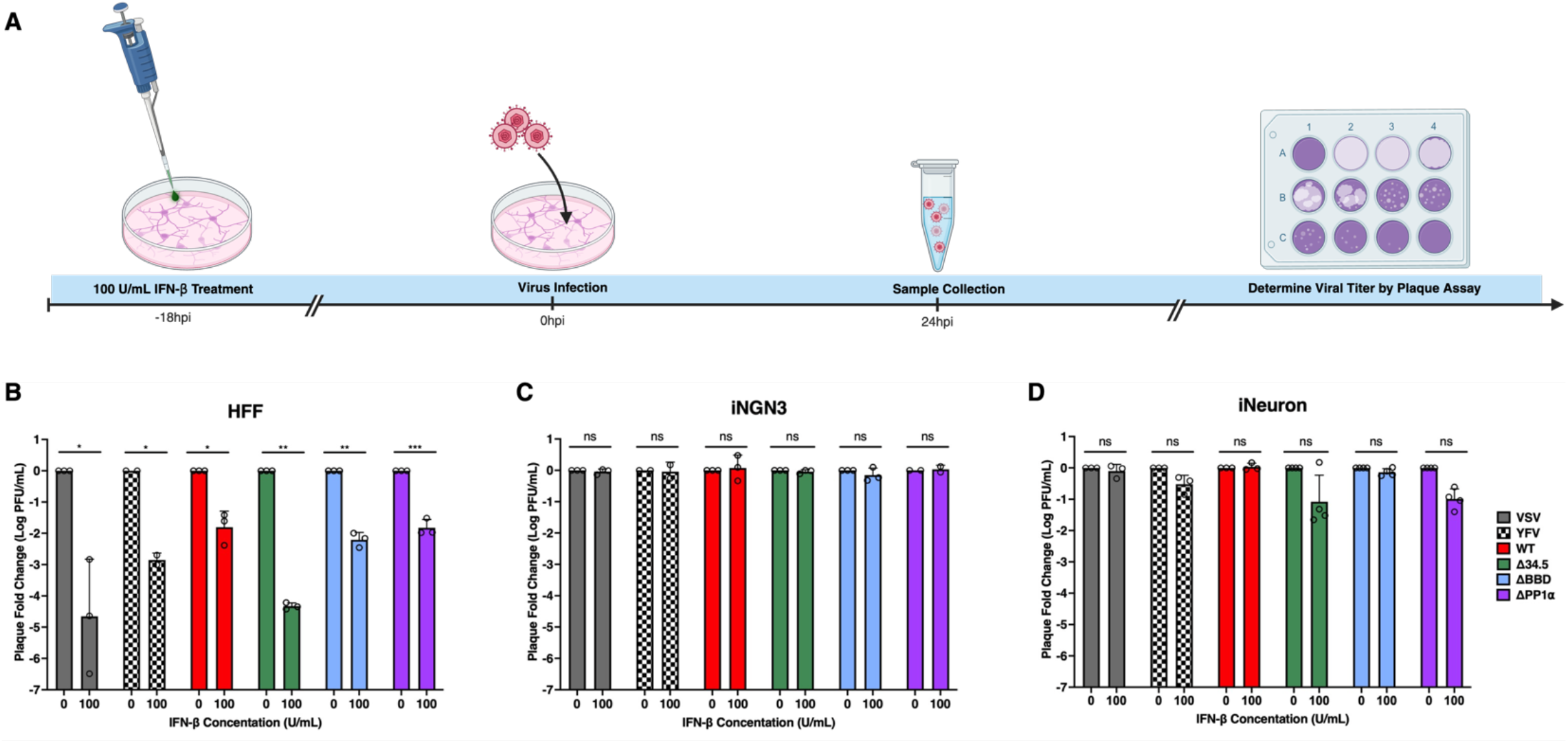
The PP1α domain of ICP34.5 mediates HSV-1 resistance to interferon. (**A**) Timeline of IFN-β treatment for experiments shown on panels **B**-**D**. Cells were treated with IFN-β for 18 hours prior to infection and collected 24 hours post infection (HPI). (**B**) Graph showing fold change of viral titers from HFFs infected with VSV, YFV, or HSV-1 recombinant viruses relative to untreated controls. (**C**) Graph showing fold change of viral titers from iNGN3 cells infected with VSV, YFV, or HSV-1 recombinant viruses relative to untreated controls. (**D**) Graph showing fold change of viral titers from iNeurons infected with VSV, YFV, or HSV-1 recombinant viruses relative to untreated controls. Each bar represents the results from mean titer of three wells repeated in three independent experiments, and displayed as the geometric mean with the error bars representing the SD. Data analyzed by Welch’s t-test. ns = not significant; *, p ≤ 0.05; **, p ≤ 0.01; ***, p ≤ 0.001.

This lack of response to type I IFNs by iPSCs led us to consider whether there is a similar deficit in iNeurons. IRF3 is activated in response to IFNs and infection[46,47], so we compared IRF3 translocation to the nucleus in iNeurons in response to IFN-β or HSV-1 as a measure of immune activation. As expected, there was an increase in nuclear IRF3 in response to IFN-β, Δ34.5, and ΔPP1α infection, but a decrease in IRF3 expression following WT infection (Supplemental 3A and 3B). Overall this demonstrated that in contrast to iPSCs, an IFN-dependent immune response can be induced in iNeurons. We next assessed the impact of IFN pretreatment of iNeurons on viral replication. IFN-β pre-treatment of iNeurons did not alter VSV replication, unlike our observation in HFFs (Figure 5D). YFV replication showed a modest yet reproducible decrease (∼3-fold) to IFN-β pre-treatment, but WT and ΔBBD HSV-1 replication in iNeurons, however, were not altered by IFN pre-treatment. This was similar to iNGN3 cells, suggesting that the BBD does not play a role in mediating IFN-resistance. In contrast, Δ34.5 replication was reduced by ∼12-fold by IFN (Figure 5D) and comparable to Δ34.5, replication of ΔPP1α was reduced ∼10-fold following IFN-β-pre-treatment in iNeurons. This indicates a role for the PP1α interaction domain of ICP34.5 in modulating the albeit limited IFN response pathway in iNeurons.

### IRF3/IRF7-mediated neuronal immunity maintains HSV-1 latency in mouse TG neurons

Our results using human iNeurons were consistent with the hypothesis that innate immune regulation by ICP34.5 not only modulates lytic infection, but also latency. To corroborate this finding in a different system, we used neurons isolated from the trigeminal ganglia (TG) of wild type adult or IRF3/IRF7 double knock-out C57BL/6 mice (Figure 6A).

**Figure 6:**
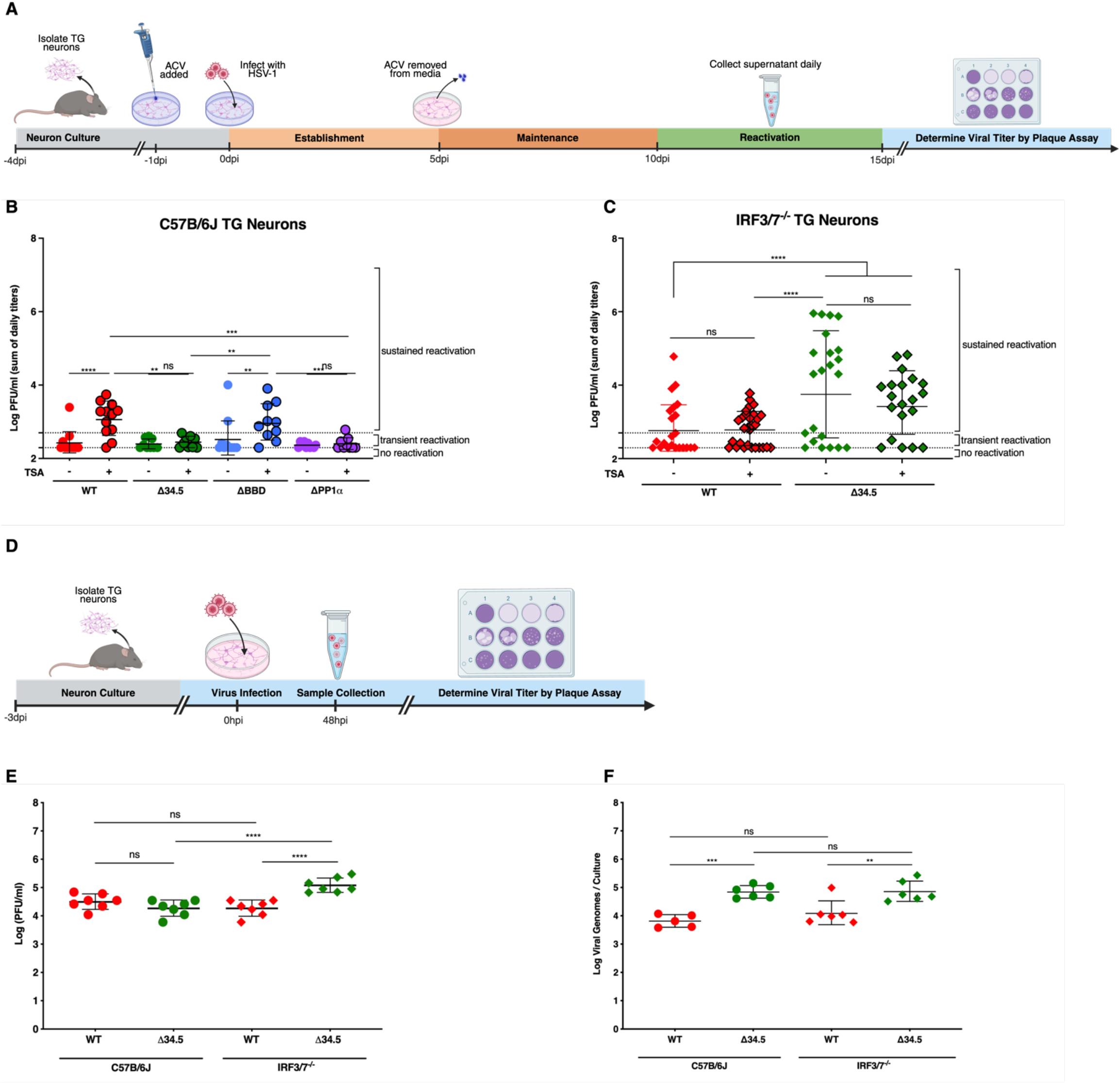
ICP34.5 counters neuronal IRF-3/7 mediated innate immunity to stimulate HSV-1 reactivation. (**A**) Timeline of establishment and maintenance of latency in mouse TG neurons. Accumulated viral titers derived from C57B/6J (**B**) or IRF3/7^-/-^ (**C**) primary murine neurons latently infected with HSV-1 recombinant viruses. Each datapoint represents the sum of viral titers detected in a single neuronal culture from 10-15DPI, and TSA-treated datapoints are shown with a black outline. The dotted line represents the limit of detection and the threshold for sustained reactivation. (**D**) Timeline of infection of TG neurons used in panel **E**. (**E**) Graph showing viral titers at 48 hours post infection (HPI) of C57B/6J (circle) and IRF-3/7^-/-^ (diamond) TG neurons infected with HSV-1 recombinant viruses (WT=red, Δ34.5=green) at an MOI of 20. (**F**) Graph showing HSV-1 genome copy number from C57B/6J and IRF-3/7^-/-^ neuronal cultures on day 5 of the latency reactivation assay. Individual circles and diamonds represent a single neuronal culture. Data represents at least 2 independent experiments per group, each with at least 3 cultures per group. Error bars represent geometric mean and standard deviation (SD). Statistical significance was determined by a two-way ANOVA (**B** and **C**) or one-way AVOVA (**E** and **F**) with Šídák posttest. ns = not significant; **, p ≤ 0.01; ***, p ≤ 0.001; ****, p ≤ 0.0001.

Briefly, excised and dissociated TG neuron cultures were established for 3 days, then pre-treated prior to HSV-1 infection with ACV for 1 day. TG neurons were infected the next day and establishment, maintenance, and reactivation of latency was measured. In TG neurons latently infected with ΔBBD, Δ34.5, or ΔPP1α there was no difference in spontaneous reactivation between any of these viruses and WT strain 17 (Figure 6B). This is in contrast to iNeurons where Δ34.5 or ΔBBD exhibited higher spontaneous reactivation (Figure 2G). Induced reactivation with TSA in TG neurons, however, revealed a different pattern, with WT and ΔBBD viruses reactivating upon TSA treatment. In contrast, the Δ34.5 and ΔPP1α viruses were unable to reactivate (Figure 6B). These data are consistent with the hypothesis that ICP34.5, through the activity of the PP1α domain, modulates reactivation through dephosphorylation of eIF2α, thus mitigating IFN-driven antiviral pathways.

Previous data demonstrated that ICP34.5 modulates type I IFN signaling through the regulation of IRF3 phosphorylation[40]. Considered together with the current data, we therefore hypothesized that regulation of IRF-dependent innate immune pathways by ICP34.5 is required for reactivation. To test this, TG neurons were prepared from mice lacking both IRF3 and IRF7 (IRF3/7^-/-^) and used for latency-reactivation experiments. IRF3 and IRF7 mediate signaling from multiple PRRs that sense HSV nucleic acids, and therefore likely facilitate detection of HSV DNA replication and reactivation events[48,49]. Consistent with our hypothesis, IRF3/7 ^-/-^ neurons infected with either WT or Δ34.5 showed sustained and spontaneous reactivation regardless of TSA treatment (Figure 6C). The robust expression of LAT and near-undetectable levels of ICP27 on day 5 postinfection suggested that a quiescent infection with latency-like gene expression was established at this time point (Supplemental Figure 4A and 4B). Together, this suggested that the IRF3/7-dependent pathways prevent reactivation of Δ34.5, and suggests that IRF3/7 is broadly instrumental in the suppression of viral reactivation. To verify that these reactivation phenotypes were not due to differences in the intrinsic abilities of these viruses to replicate in WT versus IRF3/7 ^-/-^ TG cells, we measured viral replication in these neurons (Figure 6D). As expected, WT HSV-1 and Δ34.5 had comparable viral titers in C57B/6J (B6) neurons (Figure 6E)[39]. IRF3/7^-/-^ neurons supported replication of WT HSV-1 to equivalent titers as observed in B6 neurons, while Δ34.5 grew to significantly higher titers in these cells. This raised the possibility that the discrepancy in viral replication between B6 and IRF3/7 ^-/-^ neurons during lytic Δ34.5 infection can impact the extent to which latency is established. We used qPCR to quantify HSV-1 genomes in infected TG neurons on day 5 of the latency assay, coincident with the end of establishment (Figure 6F). Mirroring the viral titer results in Figure 6E, comparable genome copy numbers of WT HSV-1 were found in B6 and IRF3/7 ^-/-^ neurons (Figure 6F). In contrast, Δ34.5 genome copy was elevated in comparison to WT HSV-1 in both B6 and IRF3/7 ^-/-^ neurons. The immunodeficiency of the IRF3/7 ^-/-^ neurons combined with this elevated copy number highlights both the multifactorial nature of latency/reactivation as well as the complexity of interpretation of data from these model systems.

## Discussion

While HSV-1 has been infecting humans for more than 5,000 years[50], there are still no vaccines or antiviral treatments that can completely break the cycle of HSV latency and reactivation. This is partly due to the lack of simple, immunologically realistic, and tractable animal models in which to study the HSV lifecycle. Historically, the most common animals to study HSV are mice, rabbits, and guinea pigs, using both *in vivo* and *ex vivo* models. While these animal models have given significant insights, they fail to mimic human immune responses and pathogenesis[24].

Furthermore, studies of the molecular basis of latency are complex with *in vivo* models. Therefore, in this study we utilized a human iPSC line that is able to differentiate into mature neuron-like cells within a week[25]. The ability of these differentiated cells to support both a productive and latent HSV-1 infection makes them a potentially attractive model to study the HSV lifecycle[31].

In response to HSV infection, multiple immune response pathways are activated[51] and HSV-1 utilizes multiple mechanisms to evade them, some through the neurovirulence factor ICP34.5[52]. ICP34.5 regulates both the IFN and autophagy pathways[18] and in this study we sought to determine roles for these functions in controlling latent infections. Autophagy is a pivotal pathway for host cell homeostasis that also serves as an antiviral response[53,54]. This can be mediated through direct degradation of intracellular pathogens, activation of IFN-driven innate immune responses, or activation of the adaptive immune response through promoting antigen presentation[14]. While it may be beneficial for the virus to inhibit autophagy in order to promote acute infection, establishment and maintenance of latency may be promoted by the virus remaining susceptible to immune clearance to promote stable latency. When iNeurons and iPSCs were infected with HSV-1, autophagic flux was unaffected. Similarly, pharmacologic autophagy modulating treatments were unable to induce significant changes in autophagy in iNeurons and failed to impact production of wild-type or ICP34.5 null HSV-1, consistent with previous observations in fibroblasts[36]. These observations support the notion that autophagy is also a proviral mechanism and a shutdown of autophagy would be detrimental to HSV at certain stages of its lifecycle[55]. Furthermore, there is a strict regulation of autophagy in iNeurons and primary neurons as it is required to maintain homeostasis in these pivotal cells[38]. As the individual steps of autophagy occur selectively within specific regions of the neuron[38], it is possible that the impact of pharmacological or viral modulation is anatomically specific and only observable under very specific experimental conditions. Consistent with this idea, mouse TG neurons respond to IFN-β differentially at the axon and soma[4]. In contrast to acute infection, we demonstrated that ICP34.5 is important for maintenance of latency in iNeurons with increased levels of spontaneous reactivation.

As we had already demonstrated that autophagy is not impacting viral production *per se* in the iNeurons, this effect might be attributable to other functions of ICP34.5, such as regulating the IFN responses through inhibiting PKR, STING, and IRF3 phosphorylation[56,57]. These IFN-β -driven antiviral responses stimulate innate immunity and inhibit viral spread[51,58] and mice strains deficient in type I IFN responses experience generalized infections and increased mortality[59,60]. This is also exhibited in humans that are deficient in the TLR3 pathway that have increased prevalence of herpes simplex encephalitis (HSE) and viral dissemination[61]. HSE has also been associated with patients with IRF3 deficiencies resulting in an inadequate type I IFN response[48]. HSV has redundant pathways to target IRF3 as a way to modulate the IFN response. As previously shown, ICP34.5 can directly and indirectly regulate IRF3[40,62]. This indirect regulation of IRF3 occurs through maintenance of ICP0[40], which facilitates degradation of IRF3 and IRF7[63]. This is consistent with our data showing an increase in IRF3 expression with Δ34.5-infected iNeurons compared to WT, and could be due to a decrease in ICP0 expression. A key point, however, is that *in vivo*, or in mixed neuronal cultures, IFN is supplied exogenously to neurons by glial cells and astrocytes. Consistent with this, when iNeurons were pretreated with IFN-β we saw small reductions in viral titers for Δ34.5 and ΔPP1α consistent with previous data in mouse TG and SCG neurons[64]. In contrast, iNGN3 cells showed no responses to IFN, consistent with previous data[45]. Notably, the reduction in viral titers were large (2-4 logs) in HFF cells, with Δ34.5 exhibiting equivalent susceptibility to VSV, the prototypic IFN-sensitive virus. This underscores the role of ICP34.5 in mediating HSV resistance to IFN as well as differences in the immune protective environment between fibroblasts and neurons.

Although ICP34.5 was required for stable latency in iNeurons, ICP34.5 was critical for robust reactivation in primary mouse neurons as seen previously[65]. Several factors could impact this differential role for ICP34.5 in these different cells. One simple explanation is the difference in species, mouse versus human cells. More likely, however, is that iNeurons consist of a homogenous culture, while mouse TG neuron cultures are heterogenous, containing glia, astrocytes, and other cell types. These supporting cells could be providing additional IFN and growth factors that suppress virus reactivation in the absence of ICP34.5, and promote cell survival. In both cell types, however, the role of ICP34.5 in the latency process appears to be more impactful on countering the IFN response than countering autophagy. Furthermore, our data with mouse IRF3/7^-/-^ TG neurons show that while the IFN response is dispensable for latency establishment, consistent with previous data[66], it is essential for controlling spontaneous reactivation.

The ability to have a host-specific cell line that is high-throughput and genetically editable to study latency provides an advantage over *in vitro* or *ex vivo* models. This has been demonstrated through the established *in vitro* latency models for other herpesviruses, such as Epstein-Barr virus (EBV) and Cytomegalovirus (CMV), that have advanced our knowledge of their molecular bases of latency[67,68]. The distinct lack of *in vitro* latency models for HSV-1 is a fundamental problem, and needs to be addressed to advance the field. Current studies aim to establish *in vitro* models to study HSV latency by utilizing neural crest-derived, neuroblastoma-derived, embryonic stem cells (ESCs), and induced pluripotent stem cells (iPSCs). These cells can be differentiated into human sensory neuron-like cells that allow for a high-throughput model to study HSV latency. However, these current cell lines can have limitations that prevent them from being widely utilized[24]. Most notably, while iPSC-derived neurons (such as iNeurons) allow for a homogenous culture by excluding glia cells, they have been shown to co-express different neuronal subtypes dependent on the differentiation protocol[27]. As previously shown in mouse TG neurons, neuronal subtype is important for latency establishment and maintenance[69]. Therefore, the disparity of neuronal subtype obtained using iPSC-derived neurons could impact HSV latency studies. In order to address this, future studies should aim to further optimize and validate latency cell lines, such as iNGN3, to make them more comparable with primary neurons and *in vivo* studies.

## Materials and Methods

### Cells

iNGN3 cells from Personal Genome Project Participant 1 were provided by Dr. George Church (Harvard University)[25]. iNGN3 cells were routinely cultured on hESC-Qualified Matrigel (Corning #354277) coated plates in mTeSR Plus media (StemCell) supplemented with 5µM Y-27632 Dihydrochloride (Sigma) and 1µM puromycin (InvivoGen) at 37°C and 5% CO_2_. The following day the media was replaced and maintained with mTeSR Plus media without supplements. Cells were split at 80-90% confluency using 0.22µM filtered 0.2mg EDTA (BioWhittaker) in 1X phosphate-buffered saline (PBS; HyClone).

iNGN3 differentiation into iNeurons was performed and cultured as previously described[31,32, Oh *et al*. Manuscript under review]. Cells are plated onto either 12mm coverslips or 96-well plates coated with 0.22 µM syringe filtered polyethylenimine (2.2 mg/mL; Sigma) for 16-20 hours at room temperature. Coverslips and plates were then washed three times with sterile water and coated with 3.3µg/mL laminin (Invitrogen) for 1 hour at 37°C and 5% CO_2_ prior to plating cells on the eighth day post induction.

Human foreskin fibroblasts (HFFs) from the American Type Culture Collection (ATCC) were cultured in Dulbecco’s modified Eagle’s medium (DMEM; HyClone SH30022.01) with 2% fetal bovine serum (FBS; HyClone) and 1% penicillin-streptomycin (HyClone) at 37°C and 5% CO_2_.

Trigeminal ganglion (TG) neurons were isolated from 6-to 10-week-old C57BL/6J or IRF3/7^-/-^ mice (Jackson Laboratory), seeded onto PDL/Laminin coated coverslips at a density of 3,600 neurons, and cultured at 37°C and 5% CO_2_ as previously described[70]. The anti-mitotic 5’-Fluro-2’ deoxyuridine (FUDR; 60µM; Sigma) was added for 3 days before cells were used in experiments.

### Immunofluorescence

Cells were washed once with PBS prior to fixation in 4% paraformaldehyde (PFA) for 15 min, and washed again with PBS. Cells were then permeabilized with 0.1% Triton-100X (Sigma) for 10 min and then washed with 0.05% Tween20. Following the wash, cells were treated at room temperature for an hour with 3% normal goat serum to block non-specific antigen binding (NGS; Vector Laboratories). Primary and secondary antibodies were diluted in PBS containing 1% NGS. Primary antibodies were added to cells overnight at 4°C, while secondary antibodies were added at room temperature for 30 min. Primary antibodies used were chicken anti-Tuj-1 (1:500; Millipore AB9354), mouse anti-NeuN (1:100; Millipore ABN91), rabbit anti-Ki67 (1:60; BioCare Medical CRM325), rabbit anti-IRF3 (1:100; Santa Cruz sc-9082), and the DNA stain 4′,6-diamidino-2-phenylindole (DAPI; Invitrogen 62248). Secondary antibodies used were goat anti-mouse Alexa 488, goat anti-rabbit Alexa 555, and donkey anti-chicken Alexa 647 (1:500; Invitrogen). All images were acquired on the Zeiss Axio Observer.Z1 microscope with a 40X objective. Images shown are representative of one (IRF3 translocation in supplemental Figure 3) or three (neuronal characterization in Figure 1) independent experiments. Fluorescent expression and translocation analysis was quantified with CellProfiler using DAPI as the nuclear stain.

### Mice

C57B6/J mice, purchased from Jackson laboratories, and IRF3/7^-/-^ mice in the C57BL/6J background[71] were bred-in house in the barrier facility in the Center for Comparative Medicine and Research at The Geisel School of Medicine at Dartmouth College. Mice were genotyped by PCR and all studies were carried out in strict accordance with the recommendations in the Guide for the Care and Use of Laboratory Animals of the National Institutes of Health. The protocol was approved by the Dartmouth IACUC Committee (5 June 2012, permit number leib.da.1, protocol # 00002151). No surgeries were performed, and all procedures were performed in strict accordance with local and Federal guidelines.

### Viruses

HSV-1 strain 17 mCherry, ΔBBD mCherry and Δ34.5 mCherry on the strain 17 background [64], HSV-1 strain 17 syn+ [72] (GenBank accession No. NC_001806.2), ΔPP1α [73], ΔBBD [19], Δ34.5 [36], and the vesicular stomatitis virus (VSV) [39] were made as previously described. The yellow fever virus strain 17D (YFV) was obtained from Charles Rice [74]. The HSV-1 strain 17 UL37-eGFP (17UL37-eGFP) and Δ34.5 UL37-eGFP viruses were generated using the pKUL37eGFP plasmid, which contains the enhanced GFP (eGFP) sequence at the C-terminus of UL37 [34]. This plasmid was linearized and cotransfected into Vero cells with HSV-1 strain 17 or Δ34.5 viral genomic DNA using Lipofectamine reagent (Invitrogen). Successful homologous recombination was tested by viral plaques and eGFP fluorescence. The resulting viruses were subjected to three rounds of plaque purification with screening by PCR and confirmation of mutation by DNA sequencing. Viral stocks were grown on African green monkey kidney (Vero) cells as described previously[75].

### Virus infection

HFFs were infected at a multiplicity of infection (MOI) of 0.1 for 1 hour at 37°C and 5% CO_2_ with agitation every 15 minutes, followed by aspiration of inoculum, and a brief wash with DMEM containing 2% FBS. Medium was replaced with DMEM containing 2% FBS and 1% penicillin-streptomycin.

NGN3s were infected at a MOI of 1 for 1 hour at 37°C and 5% CO_2_ with agitation every 15 minutes, followed by aspiration of inoculum, and a brief wash with mTeSR Plus. Medium was replaced with mTeSR Plus media with or without the autophagy modulating drug treatments LY294002 (20µM) or PIK-III (1µM) as noted.

iNeurons were infected at a MOI of 1 for 1 hour at 37°C and 5% CO_2_ without agitation, followed by aspiration of inoculum, and a brief wash with Neurobasal-A medium (Gibco). Medium was replaced with Neurobasal-A containing supplemental factors as described previously[31] with or without the autophagy modulating drug treatments LY294002 (20µM) or PIK-III (1µM) as noted.

TG neurons were cultured for 3 days prior to infection to allow for extension of neurites and infected at a MOI of 20 for 1 hour at 37°C and 5% CO_2_ without agitation. Inoculum was aspirated, cells washed with Neurobasal-A medium (Gibco), and the medium was replaced with Neurobasal-A containing supplemental factors as previously described[76].

For tittering, samples consisting of the combined supernatant and cell lysates were collected at the indicated times and frozen at -80°C. Viral titers were quantified by standard plaque assays conducted on Vero cells as described previously[75]. When noted, cells were treated with 100 U/mL of IFN-β (PBL Interferon Source) 18 hours prior to infection.

### Latency and reactivation assay

Latency and reactivation of mouse TG neurons and iNeurons was performed as previously described[77] with modifications. TG neurons were isolated as described above and cultured in Neurobasal-A media containing supplemental factors for three days (day -4). Acyclovir (ACV; 100 µM dissolved in 1M HCl) was added to TG neuron and iNeuron cultures one day prior to infection (day -1) to inhibit virus replication and promote establishment of latency. The neuronal cultures were infected on the following day (day 0) with HSV-1 recombinant viruses as noted at a MOI of 50 (TG neurons) or 1 (iNeurons) for 1 hour at 37°C and 5% CO_2_ in Neurobasal-A media without agitation, followed by aspiration of inoculum, and a brief wash with Neurobasal-A. Medium was replaced with Neurobasal-A containing supplemental factors and ACV. To ensure cell viability, media was replaced with fresh Neurobasal-A containing supplemental factors and ACV on day 3 and maintained for 2 additional days. On day 5, the media was replaced and cultured with Neurobasal-A containing supplemental factors without ACV. To ensure cell viability, media was replaced with fresh Neurobasal-A containing supplemental factors without ACV on day 8. On day 10, the media was replaced with Neurobasal-A containing supplemental factors with or without trichostatin A (TSA; 1.8µM; Sigma), which was replaced with Neurobasal-A containing supplemental factors the following day (day 11). A portion of the supernatant on days 8 and 10-15 were collected, replaced with Neurobasal-A containing supplemental factors, and assayed for infectious particles by plaque assay as described above.

For each neuronal culture, the limit of viral plaque detection by plaque assay was 28 PFU/mL. Transient reactivation was classified as 29-60 PFU/mL on a single day, or on non-consecutive days. Sustained reactivation was classified as >100 PFU/mL on three consecutive days. The sum of PFU detected throughout the course of the assay was taken as an area under the curve making the sum limit of viral plaque detection 196 PFU/mL, the sum threshold for sustained reactivation of 412 PFU/mL.

### qRT-PCR

Samples were harvested and RNA extracted using the Invitrogen PureLink RNA Micro Scale Kit. Gene expression was detected using the Luna Universal One-Step RT-qPCR Kit and 1ng of RNA with the following primer sets:

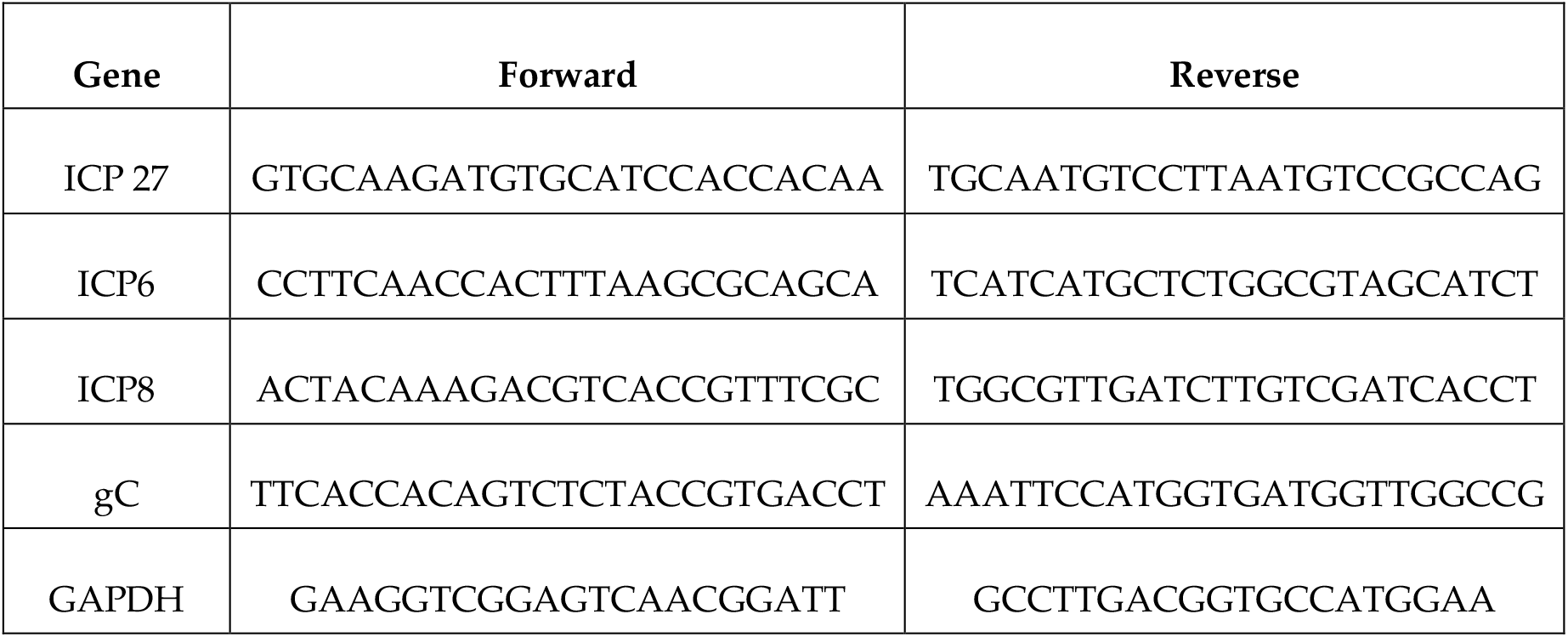

Values were obtained using the Bio-Rad CFX96 Real-Time Thermo Cycler and calculated by the 2^-ΔΔCT^ method[78] normalized to human GAPDH and then compared relative to HSV-1 strain 17 WT samples.

### p62 enzyme-linked immunosorbent assay (ELISA)

iNGN3s and iNeurons were infected with HSV-1 recombinant viruses as described above. When noted, autophagy modulating drug treatments LY294002 (20µM), PIK-III (1µM), and rapamycin (500nM; Enzo Life Sciences) were added to cultures for 24 hours.

To collect protein, media was aspirated and cells were washed once with cold PBS. Cells were lysed using 100µL of RIPA buffer containing protease and phosphatase inhibitors for 1hr at 4°C with agitation. Samples in RIPA buffer were collected into an Eppendorf tube and spun at 4°C for 5min at 14,000 rcf. Supernatant was removed and samples stored at - 80°C. Protein concentrations were determined by BCA and 50µg/mL of each sample was used with the Human p62/SQSTM1 ELISA kit as per the manufacture’s protocol (RayBiotech #ELH-SQSTM1).

### qPCR of TG neurons

DNA and RNA were harvested from latently infected murine TG neuron cultures using TRIzol™ (Thermo Fisher) and vortexed briefly to homogenize. After adding chloroform, shaking, and centrifuging, DNA was extracted from the organic layer with 4M guanidine thiocyanate, 50mM sodium citrate, and 1M Tris base and precipitated with isopropanol. RNA was purified using a MinElute PCR Purification Kit (Qiagen) and on-column RNase Free DNase Set (Qiagen).

Standards consisting of uninfected murine TG with or without the addition of *in vitro* transcribed mimics of portions of HSV-1 mRNAs were similarly prepared in TRIzol and purified in parallel. HSV-1 genome copy number was determined by qPCR using primers for viral thymidine kinase. Thymidine kinase was normalized to murine adipsin, and further normalized to the average genome load in infected neurons[79]. Reverse transcription was performed using gene specific primers for murine *GAPDH, LAT*, and *ICP27* and QuantiTect Reverse Transcription Kit (Qiagen). HSV-1 RNA expression levels for each culture were normalized to the average mGADPH levels across all the cultures. qPCR was performed with Applied Biosystems SYBR™ Green PCR Master Mix (Thermo Fisher) on a StepOnePlus machine (Applied Biosystems).

Absolute quantitation was achieved via standard curves for each assay. A thymidine kinase standard curve was established using 10-fold serial dilutions of HSV bacterial artificial chromosome (BAC) 17-49[80]. A standard curve for adipsin was similarly prepared using 5-fold serial dilutions of murine genomic material collected from a tail snip.

Primer sets used:

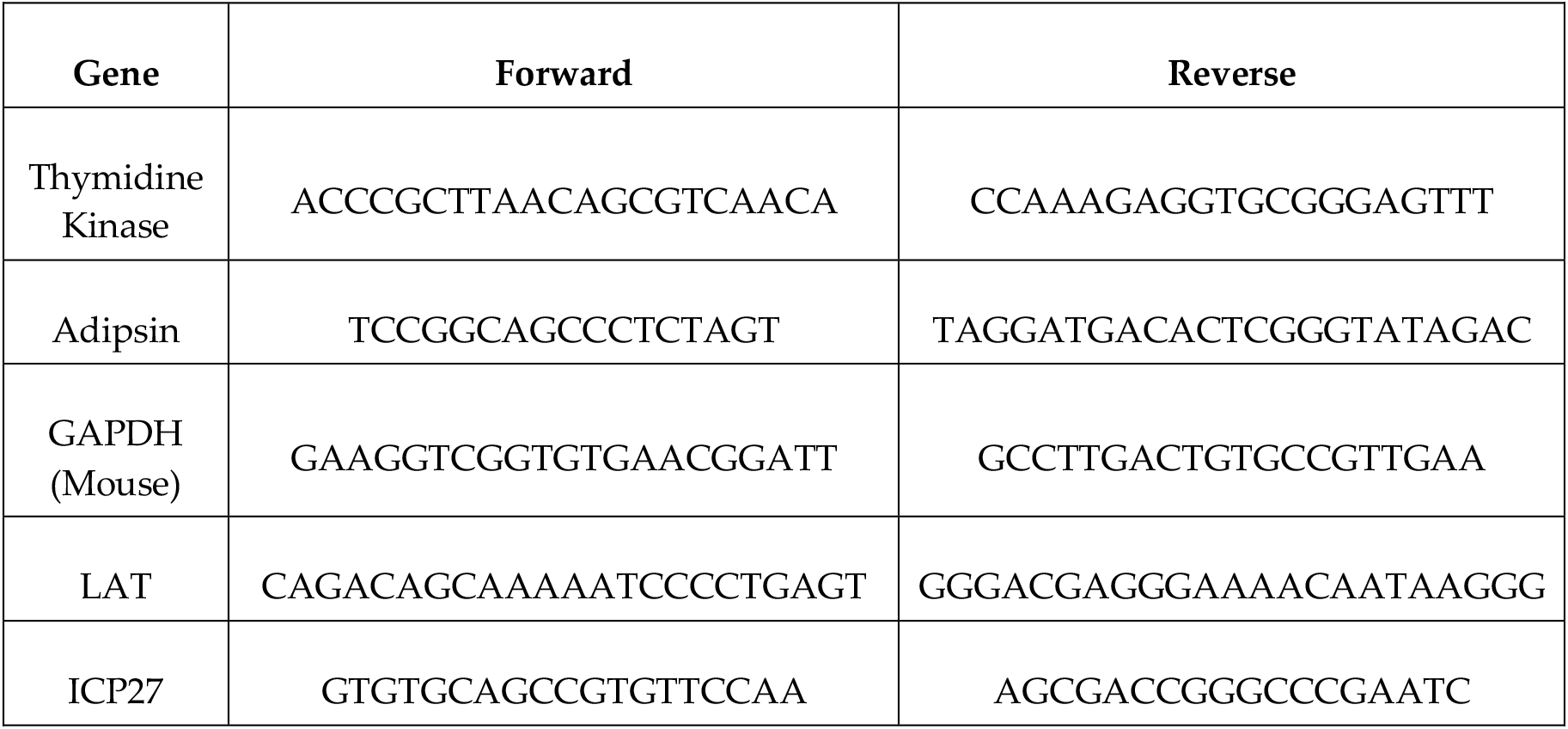

## Acknowledgements

This study was supported by National Institutes of Health (NIH) grants to D.A.L (R21 AI173941, R01 EY09083, and P01 AI098681 - also to D.M.K), and a supplement to R01 EY09083 to S.C. We also acknowledge support from the Geisel School of Medicine Immunology Program Training Grant to P.N.C, S.C., and S.K. (T32 AI007363). We also acknowledge George Church and Alex Ng for permission to the use iNGN3 cells and for technical advice.

