## Supplemental Figures for "Regulation of the innate immune response in human neurons by ICP34.5 maintains herpes simplex virus 1 latency"

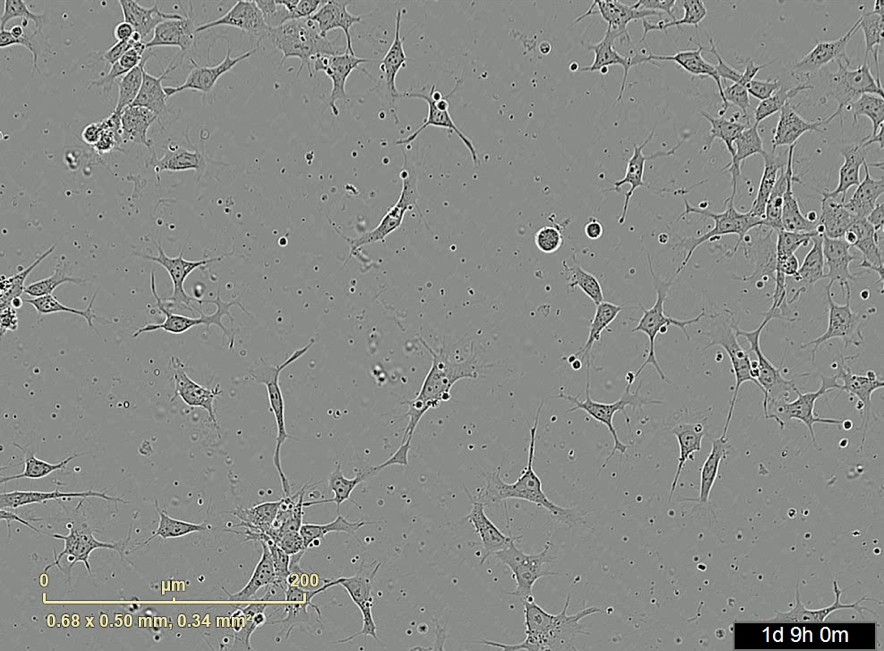

Supplemental 1: Timelapse video of differentiation of iNGN3 cells. Differentiation begins after addition of doxycycline at 1 day (1d 9h). At 3 days post induction (3d), five inhibitors (DAPT, SU-5402, CHIR99021, SB431542) are added to the media until day 8 (8d) where the media is then replaced with growth factors (BDNF, GDNF, NT3, and NGF).

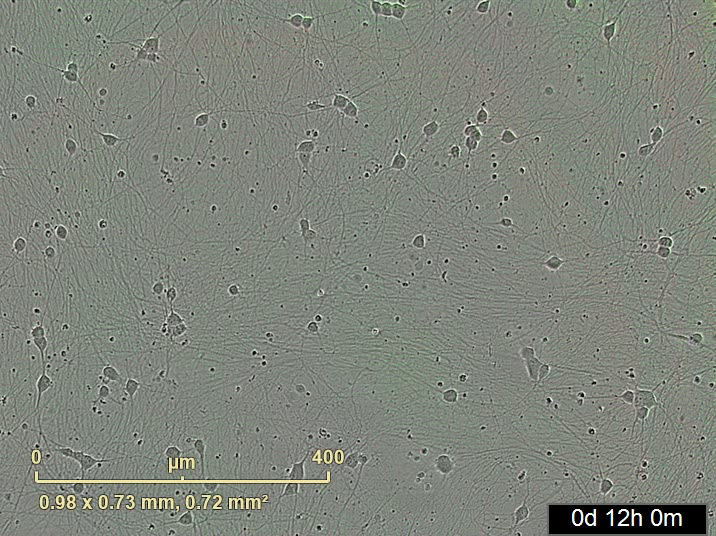

Supplemental 2: Live imaging video of fluorescent tagged HSV-1 exitng latency/quiescence in iNeurons. HSV-1 latency and reactivation was monitored through fluorescent expression from 0-15 days post infection (DPI). 100µM acyclovir (ACV) was added to the culture one day prior to infection and maintained until 5 (5d 0h 0m) days post infection (DPI). iNeurons are infected on day 0 with HSV-1 strain 17 UL37-eGFP (17UL37-eGFP) at an MOI of 1. Reactivation was induced 10DPI (10d 0h 0m) with 1.8µM trichostatin A (TSA) and TSA was removed 11DPI (11d 0h 0m).

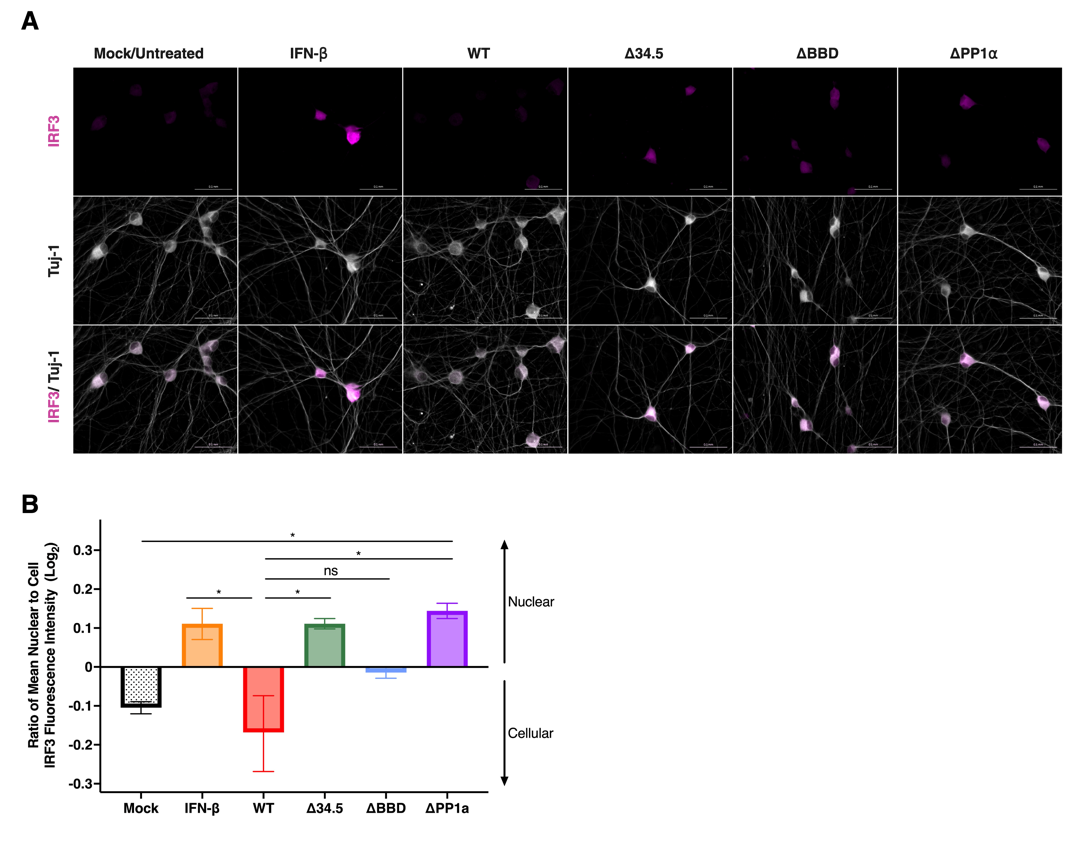

Supplemental 3: IRF3 nuclear translocation in response to IFN-β or HSV-1 infection. (A)Immunofluorescence images of IRF3 (magenta) expression in iNeurons stained with neuronal marker Tuj-1 (white). iNeurons were either treated with 100 U/mL of IFN-β for 18 hours or infected with HSV-1 recombinant viruses at an MOI of 1 for 24 hours. Scale bar = 0.1 mm. (B) Mean IRF3 fluorescence intensity per cell and mean IRF3 fluorescence intensity per nucleus was quantified by Cell Profiler. The ratio of nuclear fluorescence intensity to total cell intensity was calculated for each condition consisting of 8,000 cells. Values above the X-axis indicates a higher nuclear IRF3 signal compared to the total cellular IRF3 signal. Data analyzed by one-way ANOVA with Tukey’s posttest. ns = not significant; *, p ≤ 0.05.

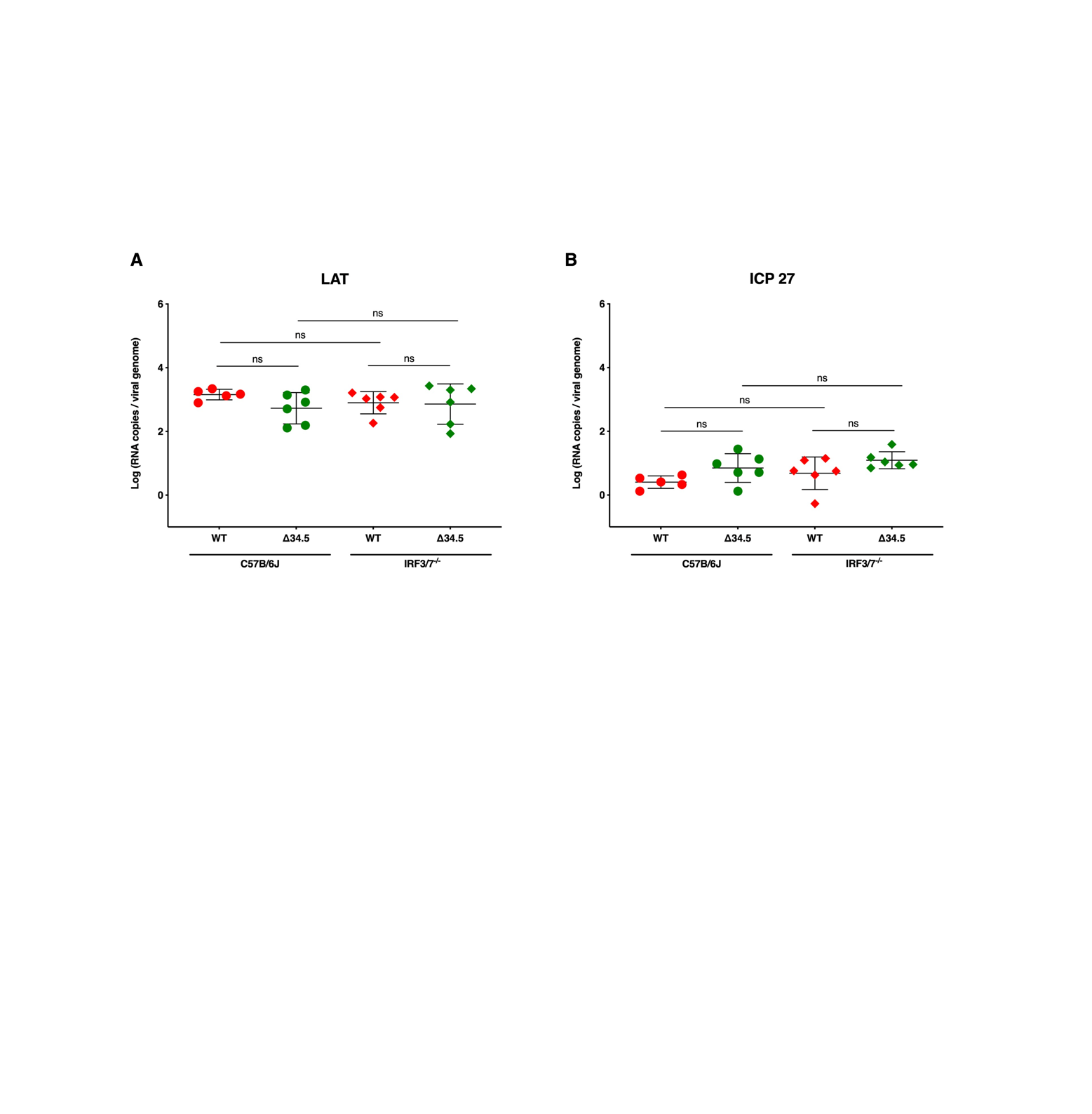

Supplemental 4: IRF3/7 is not required for establishment of quiescence in mouse neurons. (A) LAT and (B) ICP27 RNA copies were quantified by RT-qPCR and normalized to viral genome load. Individual circles and diamonds represent a single neuronal culture. Data represents 2-3 independent experiments per group, each with at least 3 cultures per group. Error bars represent standard deviation (SD). Statistical significance was determined by a one-way ANOVA with Šídák posttest. ns = not significant.
